# Comparative analysis of neuroinvasion by Japanese encephalitis virulent and vaccine strains in an *in cellulo* model of human blood-brain barrier

**DOI:** 10.1101/642033

**Authors:** Cécile Khou, Marco Aurelio Díaz-Salinas, Anaelle da Costa, Christophe Préhaud, Patricia Jeannin, Philippe V. Afonso, Marco Vignuzzi, Monique Lafon, Nathalie Pardigon

## Abstract

Japanese encephalitis virus (JEV) is the major cause of viral encephalitis in South East Asia. It has been suggested that JEV gets access to the central nervous system (CNS) as a consequence of a preceding inflammatory process which leads to the blood-brain barrier (BBB) disruption and viral neuroinvasion. However, what happens at early times of JEV contact with the BBB is poorly understood. In the present work, we evaluated the ability of both a virulent and a vaccine strain of JEV (JEV RP9 and SA14-14-2, respectively) to cross an *in cellulo* human BBB model consisting of hCMEC/D3 human endothelial cells cultivated on permeable inserts above SK-N-SH human neuroblastoma cells. Using this system, we demonstrated that both JEV RP9 and SA14-14-2 are able to cross the BBB without disrupting it at early times post-addition. Furthermore, this BBB model was able to discriminate between the virulent RP9 and the vaccine SA14-14-2 strains, as demonstrated by the presence of almost 10 times more RP9 infectious particles that crossed the BBB than SA14-14 particles at a high MOI. Besides contributing to the understanding of early events in JEV neuroinvasion, this *in cellulo* BBB model represents a suitable and useful system to study the viral determinants of JEV neuroinvasiveness and the molecular mechanisms by which this flavivirus crosses the BBB at early times of neuroinvasion.

## INTRODUCTION

*Flaviviruses* such as Japanese encephalitis virus (JEV) are arthropod-borne viruses (arbovirus) that are transmitted through the bite of an infected mosquito and may cause serious human diseases [1]. JEV is the main causative agent of viral encephalitis in South East Asia, with an annual incidence of around 68 000 cases [2]. About 30% of the cases are fatal, and half of the survivors present neurological sequelae [3]. To date, no specific treatment against JEV exists [3]. However, Japanese encephalitis is a preventable disease as vaccines have been developed: the live-attenuated JEV SA14-14-2 strain, as well as a recombinant vaccine and an inactivated one, also based on the JEV SA14-14-2 strain [4, 5]. The live-attenuated vaccine was obtained empirically after several passages of the JEV SA14 virulent strain in primary hamster kidney cells [6]. Although highly efficient, cases of post-vaccine encephalitis were also reported [7], suggesting that the vaccine strain JEV SA14-14-2 is still neurovirulent in humans.

JEV has a positive-sense RNA genome encoding a single polyprotein flanked by two untranslated regions (UTR) at its 5’ and 3’ ends. This polyprotein is co- and post-translationally cleaved into three structural proteins (capsid C, membrane prM and envelope E) involved in viral particle assembly and antigenicity and seven non-structural proteins (NS1, NS2A, NS2B, NS3, NS4A, NS4B and NS5) involved in genome replication, viral particle assembly and evasion of innate immunity [1]. Due to an error-prone NS5 polymerase that frequently introduces mutations in the viral genome during replication, a *Flavivirus* population is not clonal, but rather a mix of multiple viral genomic species (aka quasispecies) [8, 9].

JEV is a neuroinvasive and neurovirulent virus. It is associated with neuroinflammation of the central nervous system (CNS) [10], and disruption of the blood-brain barrier (BBB), as shown *in vivo* in murine and simian models [10, 11]. Expression levels of tight junction proteins involved in maintaining BBB functions such as occludin, claudin-5 and zonula occludens 1 (ZO-1) are significantly decreased in symptomatic JEV-infected mice, suggesting physical disruption of the BBB [11]. However, it seems that BBB disruption occurs after infection of the CNS cells in a mouse model of JEV-induced encephalitis [11] and that inflammatory response of infected astrocytes and pericytes plays a key role in BBB leakage [11–13], suggesting that JEV can cross the BBB before disrupting it. Indeed early studies of JEV infection of mouse brain demonstrated that the virus was transported across the cerebral endothelium by endocytosis [14]. Vesicular transport of cellular cargoes through endothelial cells is known as transcytosis [15], but it is unclear whether this mechanism also applies to the transport of JEV.

In contrast to virulent JEV strains as RP9, the vaccine strain SA14-14-2 was shown to be essentially non-neuroinvasive and non-neurovirulent in weanling ICR mice, but is still highly neurovirulent in neonates [16]. JEV SA14-14-2 genome displays 57 nucleotide differences positioned along the genome when compared to the parental strain SA14, leading to 25 amino-acid substitutions [16]. Mutations in the E protein seem to attenuate JEV neurovirulence [17, 18], while mutations in the 5’ UTR, capsid C and NS1-NS2A protein coding regions have been found to attenuate JEV neuroinvasiveness in a mouse model [18–20]. Despite the identification of these attenuating mutations, the specific amino-acids contributing to the attenuation of JEV SA14-14-2 are unknown.

Encephalitis incidents have occurred after vaccination with the SA14-14-2 JEV strain, but no virus could be recovered from them [7]. Whether these neurological adverse events originated from virus reversion to a virulent phenotype, a specific viral neuroinvasive and neurovirulent sub-population or from host determinants is still unknown [7]. In any case, the JEV vaccine strain, although much less neurovirulent and neuroinvasive than its parental counterpart, must have crossed the BBB in order to reach the CNS and initiate encephalitis.

The BBB is the physical and physiological barrier between the brain and the blood compartments in vertebrates, and it is comprised of a network of different cell types including the brain microvascular endothelium along with pericytes, astrocytes, microglia and the basement membrane [21]. Many BBB models have been developed in order to facilitate studies on the biology and pathophysiology of its diverse components, as well as to evaluate drug transport to the brain [22]. The brain microvascular endothelial cell line hCMEC/D3 exhibits a stable growth and endothelial marker characteristics that makes it suitable to form a reproducible and easy-to-grow BBB *in cellulo*. hCMEC/D3 monolayer displays good restricted permeability to paracellular tracers and retains most of the transporters and receptors present on *in vivo* BBB [23]. Accordingly, hCMEC/D3 cells have been used to investigate host-pathogen interactions with human pathogens that affect the CNS [24, 25].

In the present study, we evaluated the ability of both a virulent and a vaccine strain of JEV (JEV RP9 and SA14-14-2, respectively) to cross an *in cellulo* human BBB model consisting of hCMEC/D3 human endothelial cells cultivated on permeable supports above SK-N-SH human neuroblastoma cells. Using this system, we demonstrated that both JEV RP9 and SA14-14-2 strains are able to cross the BBB without disrupting it at early times post-addition. More importantly this BBB model is discriminant as about 10 times more RP9 than SA14-14 infectious particles may cross the barrier at a high MOI. Besides contributing to the understanding of early events in JEV neuroinvasion, this *in cellulo* BBB model represents a useful tool to examine the viral determinants of JEV neuroinvasiveness and the molecular mechanisms by which this flavivirus cross the BBB.

## MATERIAL AND METHODS

### Cell lines and JEV strains

Human endothelial cells hCMEC/D3 [23], were maintained at 37°C on rat collagen diluted at 100µg/mL in water (Cultrex; catalog no. 3443-100-01) in EndoGro medium (Merck Millipore; catalog no. SCME004) supplemented with 5% fetal bovine serum (FBS) and 10mM HEPES buffer (Sigma-Aldrich; catalog no. 83264). hCMEC/D3 cells can form tight junctions when cultured for 6 days at 37°C. Human neuroblastoma cells SK-N-SH (ATCC HTB-11) were maintained at 37°C in Dulbecco modified Eagle medium (DMEM) supplemented with 10% heat-inactivated FBS. *Cercopithecus aethiops* monkey kidney Vero cells were maintained at 37°C in DMEM supplemented with 5% heat-inactivated FBS. *Aedes albopictus* mosquito cells C6/36 were maintained at 28°C in Leibovitz medium (L15) supplemented with 10% heat-inactivated FBS.

A molecular cDNA clone of JEV genotype 3 strain RP9 was kindly provided by Dr. Yi-Ling Lin [26]. This plasmid was modified in our laboratory as previously described [27]. To produce infectious virus, the molecular clone (pBR322(CMV)-JEV-RP9) was transfected into C6/36 cells using Lipofectamine 2000 (Life Technologies; catalog no. 11668-019). Once a cytopathic effect was visible, viral supernatant was collected and used to infect C6/36 cells. As hCMEC/D3 monolayer is very sensitive to any change of medium, we used viruses produced from cells grown in the same medium as the one used to grow endothelial cells (EndoGro medium). A CD.JEVAX® (JEV SA14-14-2) vaccine vial was kindly provided by Dr. Philippe Dussart (Institut Pasteur of Phnom Penh, Cambodia). The vaccine was reconstituted with 500µL of DMEM. 250µL of reconstituted vaccine were used to infect Vero cells for 7 days. Viral supernatants were collected and used to infect C6/36 cells cultivated in EndoGro medium supplemented with 2% FBS. Both JEV RP9 and SA14-14-2 viral supernatant stocks were collected 3 days after infection and the infectious titer was determined in Vero cells through a focus-forming assay (see below).

### Antibodies

Mouse hybridomas producing the monoclonal antibody 4G2 anti-*Flavivirus* E protein were purchased from the ATCC (catalog no. HB-112), and a highly-purified antibody preparation was produced by RD Biotech (Besançon, France). Mouse monoclonal antibody anti-JEV NS5 was kindly provided by Dr. Yoshiharu Matsura [28]. Horseradish peroxidase (HRP)-conjugated goat anti-mouse IgG antibody was obtained from Bio-Rad Laboratories (catalog no. 170-6516). Alexa Fluor 488-conjugated goat anti-mouse IgG antibody was obtained from Jackson ImmunoResearch (catalog no. 115-545-003).

### Evaluation of JEV neuroinvasive capacity

5.10^4^ hCMEC/D3 cells were seeded on 12-well Transwell^®^ permeable inserts (Corning; catalog no. 3460) in EndoGro medium supplemented with 5% FBS for 5 days. 2.10^5^ SK-N-SH cells were seeded in 12-well tissue culture plates in EndoGro supplemented with 2% FBS. Permeable inserts containing hCMEC/D3 cells were then transferred in these culture plates and medium was replaced by EndoGro medium supplemented with 2% FBS. Aliquots of virus were diluted the next day in 50µL of EndoGro medium supplemented with 2% FBS, heated at 37°C and then added to the cells. Cells were incubated at 37°C until collection.

### Focus-forming assay (FFA)

Vero cells were seeded in 24-well plates. Ten-fold dilutions of virus samples were prepared in DMEM and 200µL of each dilution was added to the cells. The plates were incubated for 1h at 37°C. Unabsorbed virus was removed and 800µL of DMEM supplemented with 0.8% carboxymethyl cellulose (CMC), 5 mM HEPES buffer, 36 mM sodium bicarbonate, and 2% FBS were added to each well, followed by incubation at 37°C for 48h for JEV RP9 or for 72h for JEV SA14-14-2. The CMC overlay was aspirated, and the cells were washed with PBS and fixed with 4% paraformaldehyde for 20 min, followed by permeabilization with 0.1% Triton X-100 for 5 min. After permeabilization, the cells were washed with PBS and incubated for 1h at room temperature with anti-E antibody (4G2), followed by incubation with HRP-conjugated anti-mouse IgG antibody. The assays were developed with the Vector VIP peroxidase substrate kit (Vector Laboratories; catalog no. SK-4600) according to the manufacturer’s instructions. The viral titers were expressed in focus-forming units (FFU) per milliliter.

### Lucifer Yellow (LY) permeability assays

LY dye migration through the BBB monolayers was performed as previously described [24]. Briefly, Transwell^®^ inserts containing hCMEC/D3 monolayers were transferred in culture wells containing 1.5 mL of Hanks’ Buffered Salt Solution (HBSS) supplemented with 10 mM of HEPES buffer, 1 mM of sodium pyruvate and 50µM of LY (Sigma-Aldrich; catalog no. L0144). The culture medium inside the Transwell^®^ inserts was replaced with 500µL of HBSS buffer. Cells were incubated at 37°C for 10 min. Permeable inserts were then transferred in culture well containing 1.5 mL of HBSS buffer and incubated at 37°C for 15 min. They were then transferred in culture well containing 1.5 mL of HBSS buffer and incubated at 37°C for 20 min. Concentrations of LY in the wells were determined using a fluorescent spectrophotometer (Berthold, TriStar^2^ LB 942). The emission at 535 nm was measured with an excitation light at 485 nm. The endothelial permeability coefficient of LY was calculated in centimeters/min (cm/min), as previously described [29].

### Virus infections

10^5^ hCMEC/D3 cells were seeded on coverslips in 24-well tissue culture plates in EndoGro medium supplemented with 5% FBS. After 5 days, cell medium was replaced with 1 mL of EndoGro medium supplemented with 2% FBS. 10^5^ SK-N-SH cells were seeded on coverslips in 24-well tissue culture plates in DMEM supplemented with 2% FBS. Aliquots of virus were diluted in 200µL of medium and added to the cells. Plates were incubated for 1h at 37°C. Unabsorbed virus was removed and 1mL of EndoGro or DMEM supplemented with 2% FBS was added to the cells, followed by incubation at 37°C until collection.

### Immunofluorescence analysis (IFA)

All the following steps were performed at room temperature. Cells were fixed with 4% paraformaldehyde for 20 min followed by permeabilization with 0.1% Triton X-100 for 5 min. After permeabilization, the cells were washed with PBS and incubated for 5 min with PBS containing 1% BSA. The cells were then washed with PBS and incubated for 1h with anti-JEV NS5 antibody diluted at 1:200 in PBS, followed by incubation with Alexa Fluor 488-conjugated anti-mouse IgG antibody diluted at 1:500 in PBS. The coverslips were mounted with ProLong gold antifade reagent with DAPI (Life Technologies; catalog no. P36931). The slides were examined using a fluorescence microscope (EVOS FL Cell Imaging System).

### Gene expression studies

5.10^4^ hCMEC/D3 cells were seeded on 12-well Transwell® insert filters in EndoGro medium supplemented with 5% FBS for 5 days. 2.10^5^ SK-N-SH cells were seeded in 12-well tissue culture plates in EndoGro supplemented with 2% FBS. Transwell^®^ containing hCMEC/D3 cells were then transferred in these culture plates and medium was replaced by EndoGro medium supplemented with 2% FBS. Cells were incubated at 37°C. At 24h post-contact, total RNA of hCMEC/D3 cells were extracted using NucleoSpin RNA kit (Macherey-Nagel; catalog no. 740955.50) following the manufacturer’s instructions. 200 ng of total RNA were used to produce cDNA using the SuperScript II Reverse Transcriptase (Thermo Fisher; catalog no. 18064014) according to the manufacturer’s instructions. Quantitative PCR were performed on 2µL of cDNA using SYBR Green PCR Master Mix (Thermo Fisher; catalog no. 4309155) according to the manufacturer’s instructions. The CFX96 real-time PCR system (Bio-Rad) was used to measure SYBR green fluorescence with the following program: an initial PCR activation at 95°C (10 min), 40 cycles of denaturation at 95°C (15s) and annealing-extension at 60°C (1 min). Results were analyzed using the CFX Manager Software (Bio-Rad) gene expression analysis tool. GAPDH was used as the reference gene. Primers used in gene expression studies are listed in Table 1.

**Table 1.**
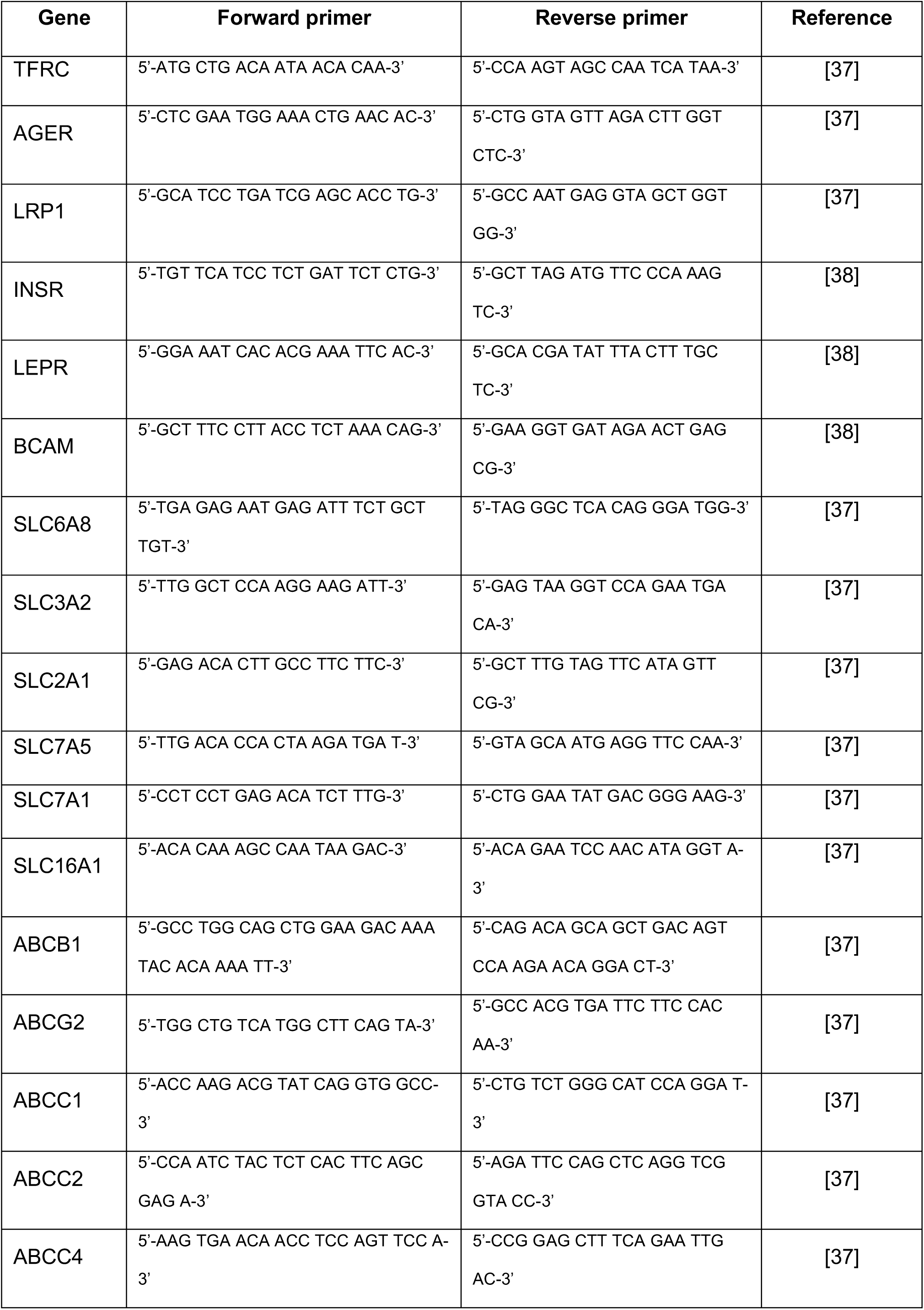
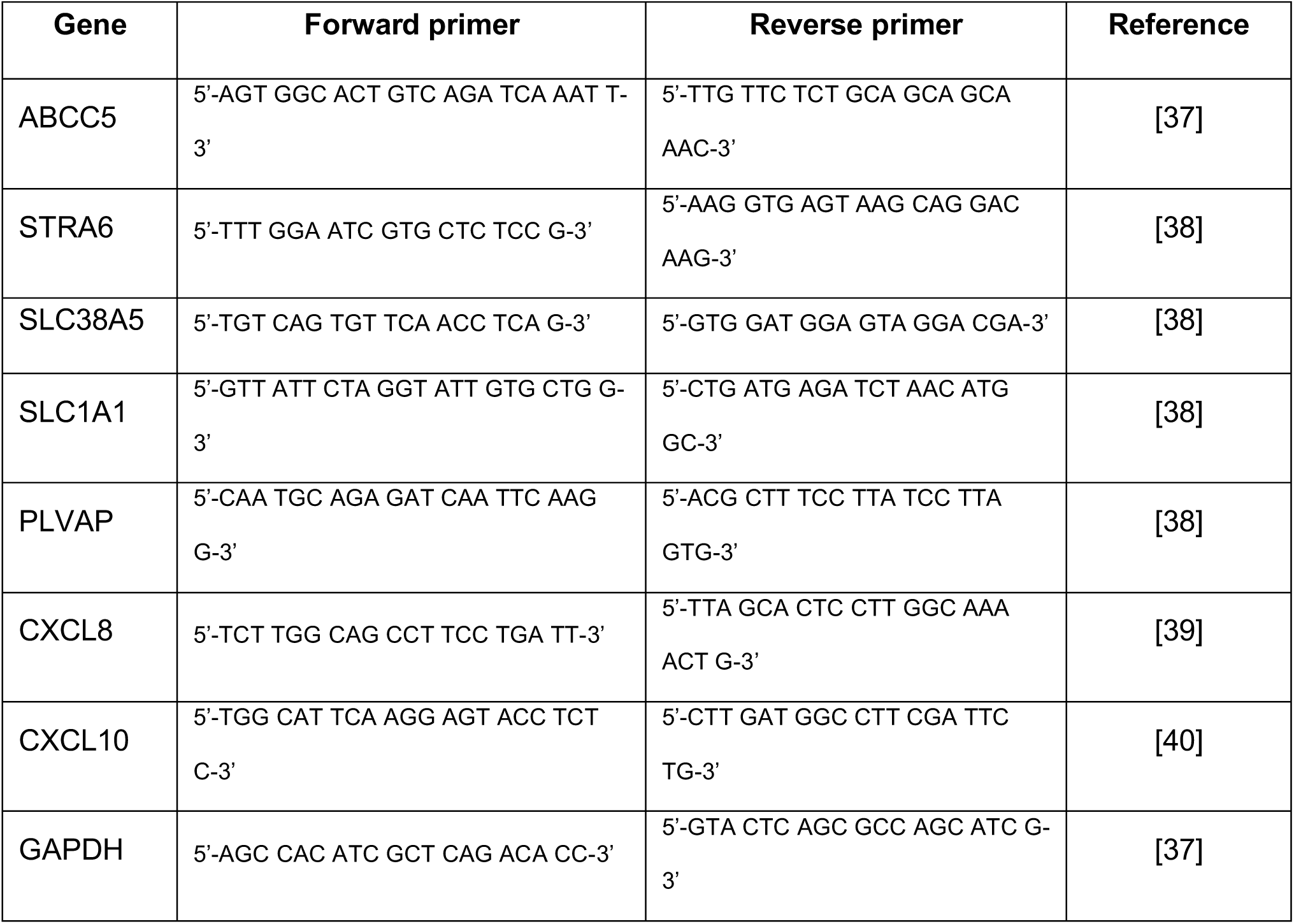
Primers used for quantification of tight junctions, receptors and transporters encoding genes.

### Quantification of JEV RNA copies number

Total RNA from JEV BBB-crossing samples was extracted using NucleoSpin®RNA kit (Macherey-Nagel; 740955.50) according to the manufacturer’s instructions. The number of JEV RNA copies present in BBB-crossing samples was determined by RT-qPCR using TaqMan® Fast Virus 1-Step Master Mix kit (Applied Biosystems®, 4444432) according to the manufacturer’s instructions. The forward and reverse primers (Sigma-Aldrich®) were 5’GAAGATGTCAACCTAGGGAGC3’ and 5’TGGCGAATTCTTCTTTAAGC3’ respectively, while [6FAM]AAGAGCCGTGGGAAAGGGAGA[BHQ1] was the probe for the assay. JEV RNA copies were calculated from a standard curve generated by amplifying known amounts of *in vitro*-transcribed RP9 NS5 gene region cloned and under SP6 promotor control. The *in vitro* transcription was performed using mMESSAGE mMACHINE™ SP6 kit (Invitrogen, Thermo Fisher Scientific, AM1340) following the manufacturer’s instructions.

### Statistical analysis

Unpaired two-tailed *t* test, Mann-Whitney test and ANOVA test corrected with Tukey method for multiple comparisons were used to compare experimental data. GraphPad Prism 7 was used for these statistical analyses.

## RESULTS

### hCMEC/D3 cell monolayers grown on permeable inserts form a BBB whose properties are not affected by SK-N-SH cells presence

A basic *in cellulo* model to study JEV neuroinvasion should consist of two main components: 1) a cell monolayer mimicking the BBB, and 2) a brain tissue-derived cell line permissive to JEV. Based on our previous work [24], we chose to use hCMEC/D3 human endothelial cells monolayers cultivated on permeable inserts and place these inserts in wells in which human neuroblastoma SK-N-SH cells were grown, in order to partly mimic the brain parenchyma. Relevant parameters of a functional BBB model, such as permeability and presence of cell transporters and receptors specific of hCMEC/D3 cells were evaluated when the endothelial cells were or not grown above SK-N-SH monolayers (Fig. 1). Permeability measurement of hCMEC/D3 monolayers through evaluation of Lucifer Yellow (LY) passage showed no significant difference whether SK-N-SH cells were present or not (Fig. 1A, + or - respectively). Moreover, the relative RNA level of genes coding for proteins involved both in cell receptors (Fig. 1B) and transporters (Fig. 1C) characteristic of endothelial barriers were similar in the two conditions, suggesting that the culture of neuroblastoma cells under the inserts on which hCMEC/D3 were grown did not disturb the endothelial cell intrinsic BBB properties and actually makes of this *in cellulo* BBB model a useful tool to study the neuroinvasion ability of JEV.

**Fig. 1.**
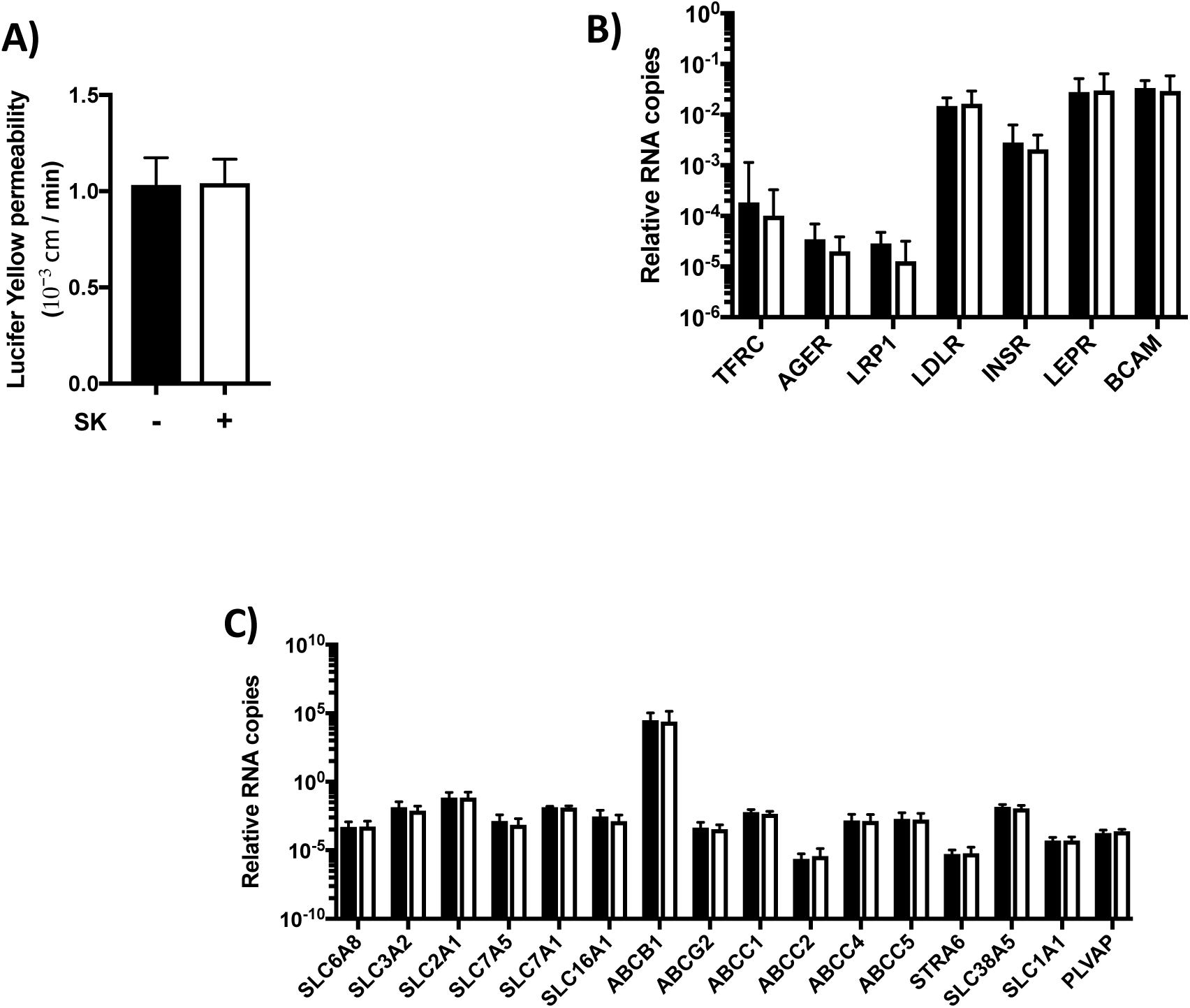
The presence of SK-N-SH cells under hCMEC/D3 BBB-forming cells does not affect the BBB properties. hCMEC/D3 were cultivated on Transwell^®^ inserts. Five days after seeding, SK-N-SH (SK) cells were cultivated or not in wells under the Transwell^®^ inserts (white and black bars respectively). **A)** Twenty-four hours after adding the SK-N-SH cells (+) or not (-), BBB permeability to LY was measured. **B)** and **C)** hCMEC/D3 BBB-forming cells total RNA was extracted and receptors (B) and transporters (C) typical of the BBB-encoding genes were quantified by RT followed by qPCR as described in Material and Methods. Graphs show the results from two independent experiments performed by duplicates.

### JEV SA14-14-2 is less replicative than JEV RP9 in SK-N-SH cells

Neuroblastoma SK-N-SH cells are susceptible to both the virulent JEV RP9 strain and the SA14-14-2 attenuated strain [27, 30]. However, a direct comparison between replication of these two JEV strains in that cell line has not been described. We thus evaluated replication of each JEV strain in SK-N-SH cells at 24 and 48 hpi (Fig. 2). As expected, both JEV strains infected the neuroblastoma-derived cell line as demonstrated by the detection of a viral antigen (NS5 protein) through immunofluorescence assays (Fig. 2A). However, the viral progeny of JEV SA14-14-2 vaccine strain produced in SK-N-SH cells at 24 and 48 hpi was significantly lower than that of JEV RP9 (1.7 and 1.2 log10 less at 24 and 48 hpi respectively, Fig. 2B), suggesting that JEV SA14-14-2 is less neurovirulent than JEV RP9 in human cell cultures.

**Fig. 2.**
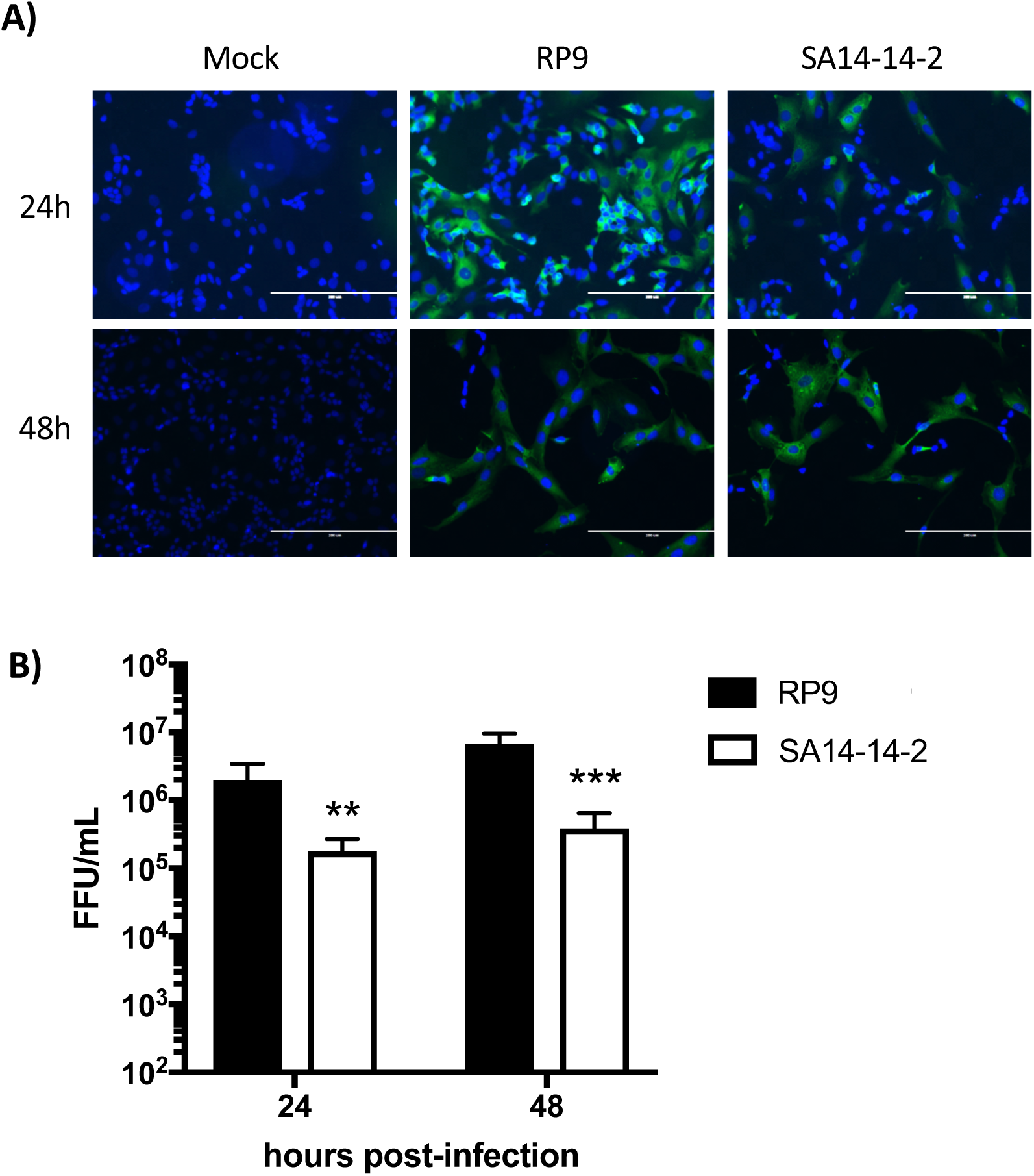
JEV RP9 is more replicative than JEV SA14-14-2 in SK-N-SH neuroblastoma cells. SK-N-SH cells were infected at MOI 0.1 for 24 or 48h by the indicated JEV strain. **A)** The infected cells were analyzed at the indicated times post-infection by immunofluorescence staining for the presence of the NS5 protein (in green). The images were taken at a x200 magnification, the cell nuclei were stained by DAPI (in blue). **B)** Supernatants of SK-N-SH cells infected by JEV RP9 (black bar) or JEV SA14-14-2 (white bar) were titrated in Vero cells. The arithmetic means ± standard deviation of three independent experiments performed in triplicate is shown. Asterisks indicate a significant difference between RP9 and SA14-14-2 in each one of the times post-infection evaluated (**, P < 0.01, ***, P < 0.001).

### Neither JEV RP9 nor JEV SA14-14-2 infects hCMEC/D3 cells after they form a BBB

In order to examine the susceptibility of our hCMEC/D3 BBB model to JEV infection, the cells were grown 6 days on coverslips to allow the BBB to form, then inoculated with either RP9 or SA14-14-2 JEV strain (Fig. 3). The presence of the NS5 viral protein as infection evidence was assessed by immunofluorescence microscopy as described in the Material and Methods section. No fluorescence signal was observed in hCMEC/D3 BBB-forming monolayer either at 24 or 48 hpi (Fig. 3A). Surprisingly, hCMEC/D3 cells could be infected by either JEV strains when they were inoculated after only one day of culture (ie not forming of a BBB), as detected through the same immunofluorescence approach (Fig. 3B). Moreover, in this condition, both JEV strains produced infectious viral progeny in hCMEC/D3, although the RP9 viral titer was significant higher by around 2 log than that observed for SA14-14-2 (Fig. 3C). These results suggest that hCMEC/D3 cells are not susceptible to JEV infection when they already have formed a barrier, but they are JEV permissive before tight junctions formation.

**Fig. 3.**
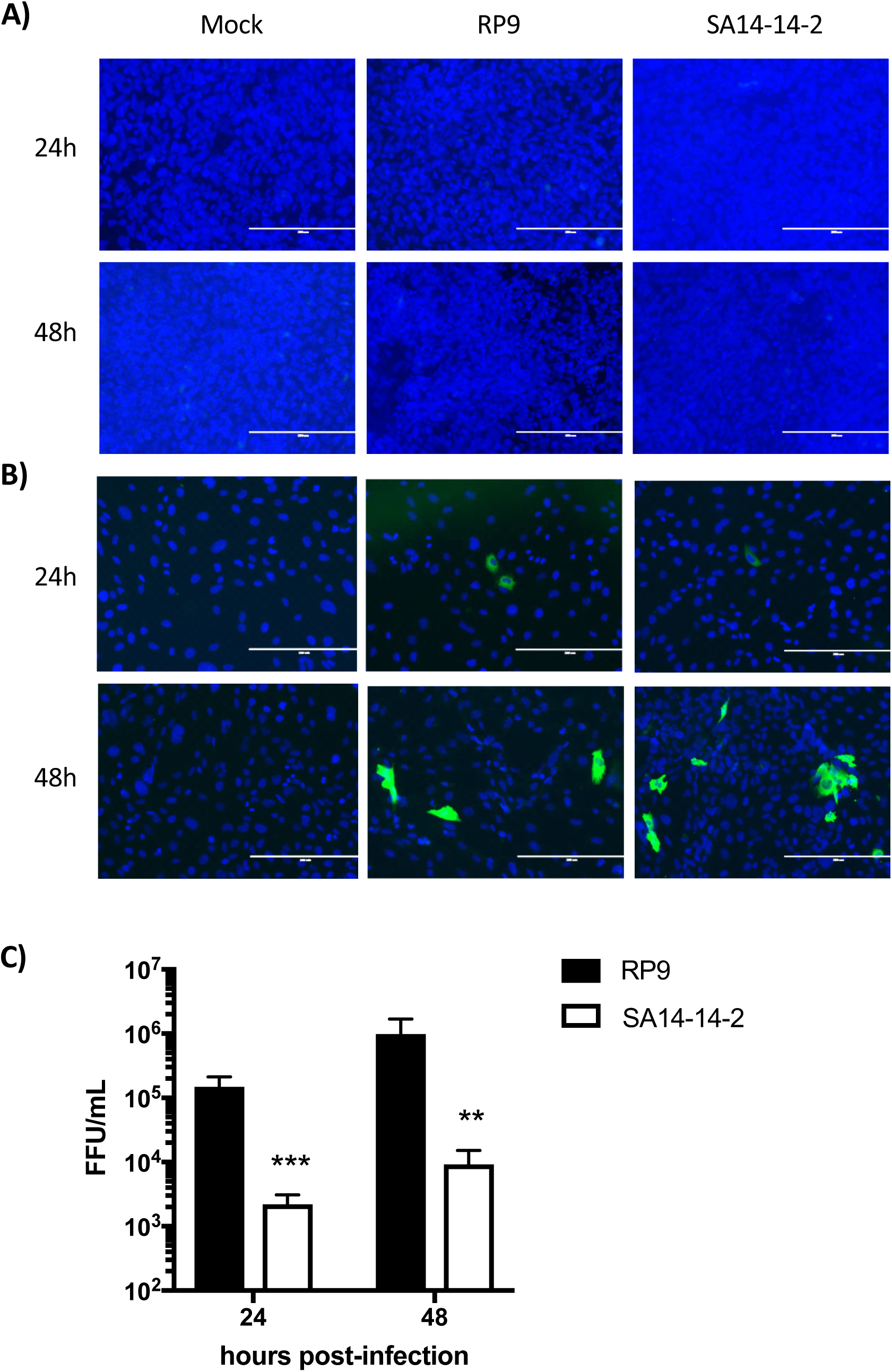
Infection of hCMEC/D3 cells by JEV strains. hCMEC/D3 were cultured on coverslips for either 6 days (**A**) or 1 day (**B**), so that they form or not a BBB respectively. Cells were then inoculated with the indicated JEV strain at MOI=0.1 and analyzed at 24 and 48 hpi by immunofluorescence staining for the presence of the NS5 protein (in green). The images were taken at a x200 magnification, the cells nuclei are stained by DAPI (in blue). **C)** Supernatants from hCMEC/D3 cells that do not form a BBB and infected by JEV RP9 (black bar) or JEV SA14-14-2 (white bar) were collected at 24 and 48 hours post-infection and their viral titer was determined as described in Material and Methods. The arithmetic means ± standard deviation of three independent experiments performed by triplicate is shown. Asterisks indicate a significant difference between RP9 and SA14-14-2 in each one of the times post-infection evaluated (**, P < 0.01, ***, P < 0.001).

### Neither JEV RP9 nor JEV SA14-14-2 disrupts the BBB when added for 6h

It has been suggested that JEV infects brain tissue cells as a consequence of a preceding inflammatory process which leads to the BBB disruption and viral neuroinvasion [31, 32]. However, the very early events of JEV BBB crossing are still poorly understood. In order to evaluate the neuroinvasive ability of JEV in our BBB model at early times post-addition, hCMEC/D3 cells cultivated on permeable inserts to form a BBB above SK-N-SH cells monolayer were exposed to either JEV RP9 or SA14-14-2 virus addition (MOI=1 or 10; Fig. 4). The permeability of the BBB at 6 hpi in the presence of the 2 different JEV strains did not show a significant difference when compared to that of the mock-infected condition (Fig. 4A), suggesting that the BBB model was not disturbed either by the JEV strains or the MOIs used.

**Fig. 4.**
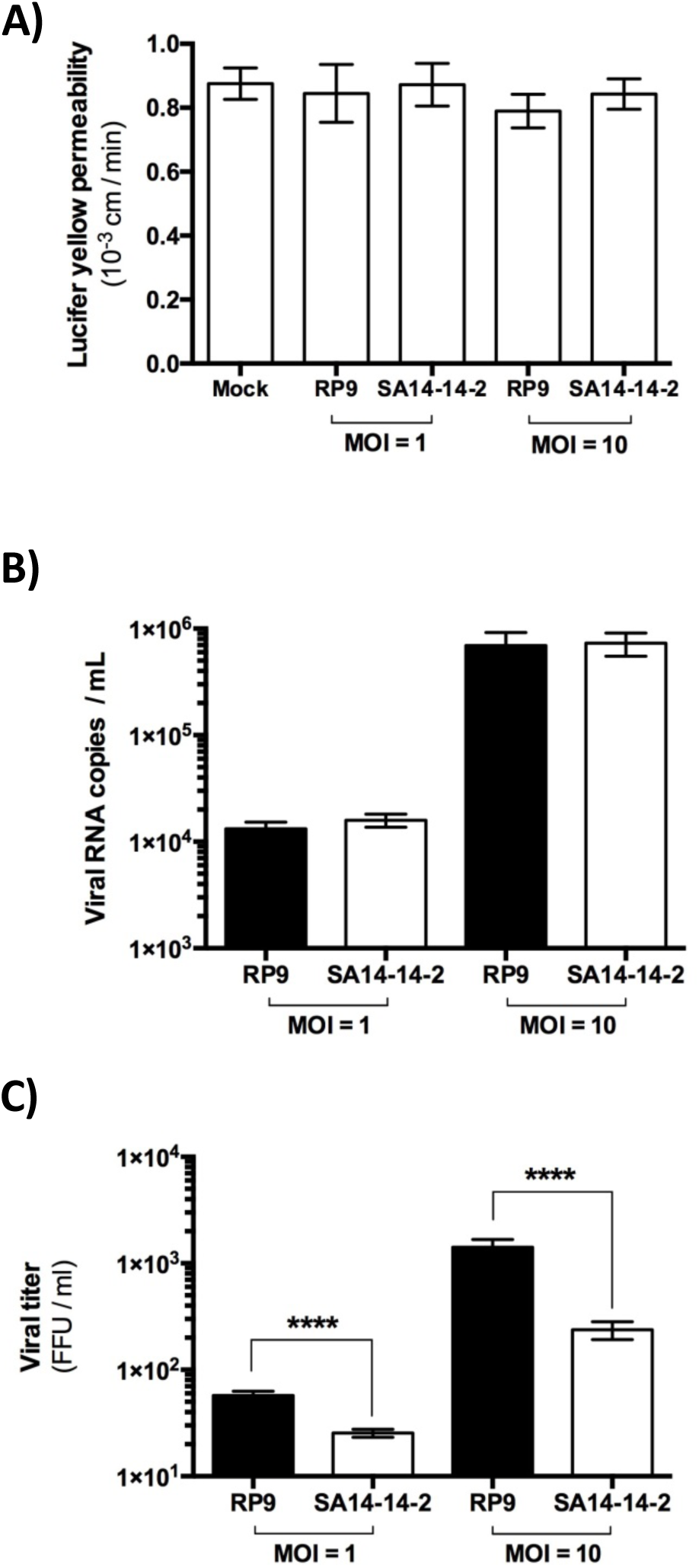
JEV RP9 and JEV SA14-14-2 may cross the *in cellulo* BBB model without disrupting it. **A)** hCMEC/D3 cells were cultivated on Transwell^®^ inserts. Five days after seeding, SK-N-SH cells were added to the wells under the Transwell^®^ insert. 24h later JEV RP9 or SA-14-14-2 was added either at MOI=1 or =10 to the BBB as indicated. Permeability to Lucifer Yellow was assayed in the hCMEC/D3 cells 6 h post-addition as described in Material and Methods. **B)** *In cellulo* BBBs were generated as indicated above and either JEV strain was added at MOI=1 or =10. After 6 h, total RNA was extracted from JEV BBB-crossing samples under the inserts and the number of JEV RNA copies was determined by RT-qPCR as described in Material and Methods. **C)** Either JEV RP9 (black bars) or SA14-14-2 (white bars) BBB-crossing samples were collected after 6 h post-addition and their viral titer was determined as described in Material and Methods. The arithmetic means ± standard deviation of at least two independent experiments performed by triplicate is shown. Asterisks indicate a significant difference between the RP9 and SA14-14-2 titers in each one of the MOIs evaluated in the BBB-crossing experiments (****, P < 0.0001).

### More JEV RP9 infectious particles may cross the *in cellulo* BBB model than JEV SA14-14-2

Since the BBB permeability was not affected by the addition of either virus, we examined the viral crossing of each strain by evaluating the quantity of viral RNA and infectious particles in the supernatants under the inserts (Fig. 4B and C). The number of viral RNA copies detected for both viruses was 1.7 log10 higher when a MOI of 10 was used in comparison to a MOI of 1 (Fig. 4B), suggesting that the higher the JEV viral load, the greater the number of viral particles crossing the BBB. Of note, there was no significant difference in the viral RNA copy number between the JEV strains for each MOI (MOI=1 or =10, Fig. 4B). However, the infectious titers of the JEV particles that crossed the BBB was surprisingly different between the RP9 and SA14-14-2 strains, as about 3 times more RP9 infectious particles where found in the supernatants under the inserts than SA14-14-2 when an MOI of 1 was used, and close to 10 times for a MOI of 10 (Fig. 4C). Calculation of the specific infectivity for JEV RP9 and SA14-14-2 strains as the ratio between the detected JEV RNA copy number per infectious focus-forming unit did not show a significant difference between the 2 viral stocks (Fig. 5A). Interestingly, the specific infectivity for the RP9 BBB-crossing samples was significantly lower than that observed for the vaccine strain SA14-14-2 with a 3 to 10 fold decrease for MOI=1 and =10 respectively (Fig. 5B). These results indicate that more JEV RP9 infectious particles may cross our BBB model than SA14-14, and demonstrate that this *in cellulo* barrier is capable of discriminating between 2 viruses with different neuroinvasive capabilities.

**Fig. 5.**
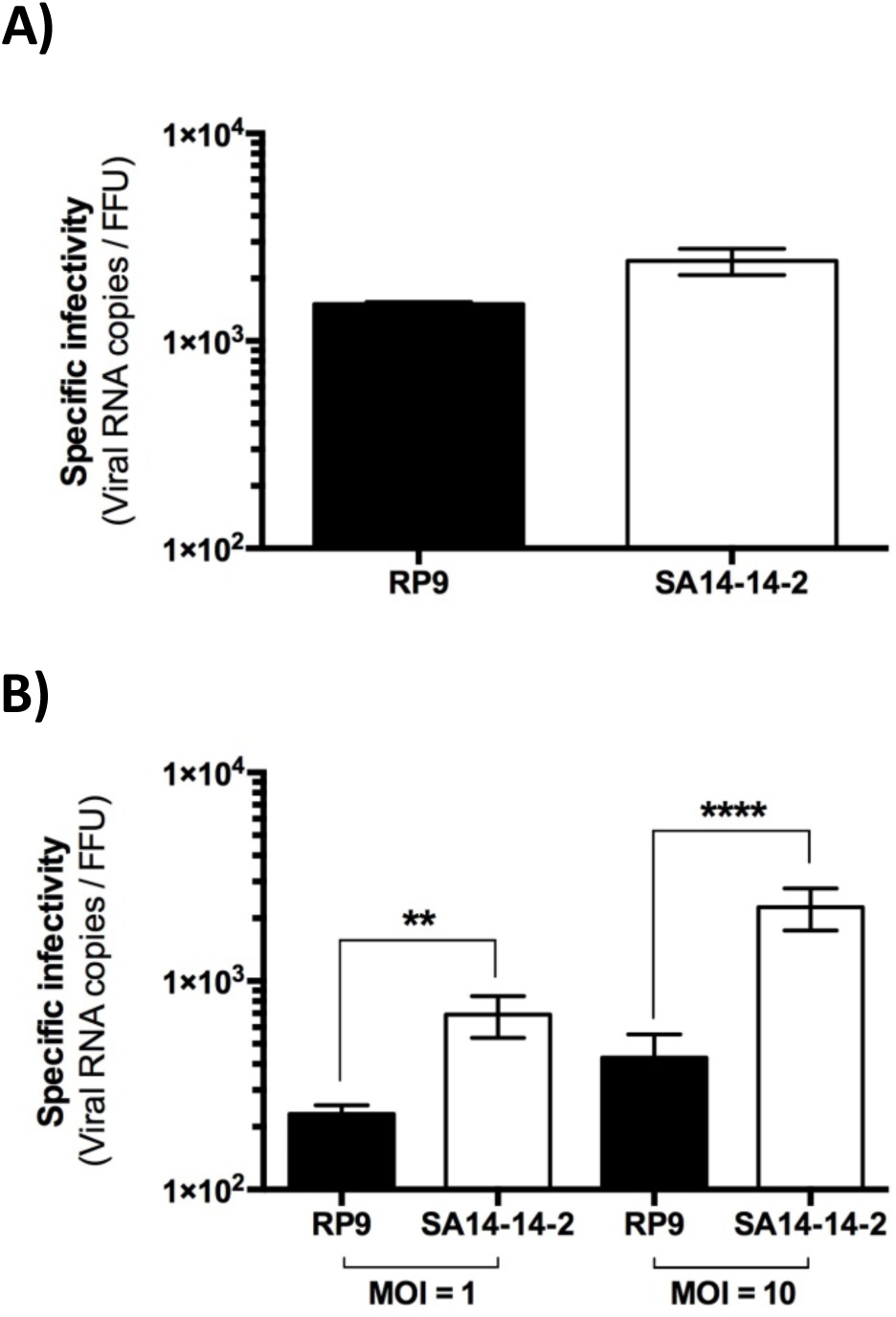
The specific infectivity of JEV RP9 is decreased after BBB-crossing. **A)** Both the number of viral RNA copies and the infectious titers for either JEV RP9 or JEV SA14-14-2 stocks used for the JEV BBB-crossing experiments were determined as described in Material and Methods. The specific infectivity of both stocks was calculated by dividing the viral RNA copies number/ml by the FFU/ml of each viral stocks. B) The specific infectivity of the JEV RP9 and SA14-14-2 BBB-crossing samples was calculated as indicated above using the data from Fig. 4B and 4C. The arithmetic means ± standard deviation of at least two independent experiments performed by triplicate is shown. Asterisks indicate a significant difference between the specific infectivity of RP9 and SA14-14-2 in each one of the MOIs evaluated in the BBB-crossing experiments (**, P < 0.01; ****, P < 0.0001).

## DISCUSSION

Several lines of research in either *in vivo* and *in vitro* systems have suggested that JEV infects brain tissue cells as a consequence of a preceding inflammatory process which would lead to the BBB disruption and viral neuroinvasion [31, 32]. While *in vivo* approaches are useful to understand the systemic viral disease, *in vitro* models are also useful because they allow studying the molecular mechanisms that govern viral pathogenesis. In this regard, previous approaches have been used to characterize JEV neuroinvasion properties at late times of infection, mainly 24 hpi or later [30, 33–35]. However, knowledge relative to the early times of JEV contact with the BBB is poor, if not null.

In this study, we have used an *in cellulo* model of a human BBB to compare JEV RP9 virulent and JEV SA14-14-2 vaccine strain ability to cross the BBB at early times post-addition. We have shown that both JEV RP9 and SA14-14-2 are able to cross the BBB without disrupting it at 6 hpi (Fig. 4). This finding is very relevant because it suggests that JEV could be able to get access to the CNS and establish a primary infection there without the preceding need of inflammatory cytokines that could lead to BBB disruption prior CNS cells viral infection as it is currently thought [31, 32].

Moreover, the fact that both JEV RP9 and SA14-14-2 strains crossed the BBB without infecting its endothelial cells, nor disrupting the barrier, also suggests that JEV is able to cross the BBB in a transcellular way through the endothelial cells or in a paracellular way between the endothelial cells. These observations are consistent with other studies conducted *in vivo* in mice and monkeys [11, 16, 36]. Observations of JEV-infected suckling mice brain by electron-microscopy suggested that JEV crosses the BBB endothelial cells by transcytosis [14]. Regardless of these observations, currently there is no published data to support this hypothesis from biochemical, genetics or functional approaches. The combination of these approaches, together with the use of our *in cellulo* BBB model and JEV strains with different neuroinvasive capabilities such as the ones used in this work would be useful to identify which cell mechanisms are “highjacked” by these pathogens to cross the BBB.

Interestingly, our specific infectivity data suggest that JEV RP9 infectious particles crossed the BBB more efficiently than JEV SA14-14-2 (Fig. 5). Comparison of the transcriptome from hCMEC/D3 cells forming a BBB to which either JEV RP9, SA14-14-2 or no virus was added for 6h showed no significant difference in the levels of gene expression (fold-change threshold of 2, data not shown). This suggests that an early cell response is not responsible for the differential BBB crossing of JEV RP9 versus JEV SA-14-14-2 particles we observed (Fig. 4C), and that it most likely stems from viral factors. It is also possible that the difference in the JEV RP9 and SA14-14-2 infectious particles ability to cross the *in cellulo* BBB relies on specific protein interactions, for example, interaction of the viral particle with a strain-specific cell surface receptor for viral entry. A full characterization of the viral particles that are able to cross the BBB including by deep-sequencing of their RNA content and examining the endothelial cells forming the BBB after contact with either virus by electron microscopy, together with uncovering specific cell receptor(s) for JEV strains could help solving these issues.

Surprisingly, we found that hCMEC/D3 were permissive to both RP9 and SA14-14-2 strains only when the BBB formation was not completed (Fig. 3B), suggesting that formation of tight junctions between these cells could make the JEV cell entry receptor(s) inaccessible to the virus. Based on our data and considering the current model of JEV neuroinvasion that suggests disruption of the BBB following CNS viral infection [11], endothelial cells from a disrupted barrier might become permissive to JEV because of better accessibility to cell entry receptor(s), and these cells, upon infection, could in turn become a new source of viral production contributing to JEV infection of the CNS.

In conclusion, our study demonstrates that both JEV RP9 and SA14-14-2 are able to cross a BBB model without disrupting it at early times post-addition and that the BBB formed by human endothelial cells represents a useful discriminant *in cellulo* model to characterize viral determinants of JEV neuroinvasiveness as well as a tool to study the molecular mechanisms by which these pathogens cross the BBB.

## ACKNOWLEDGMENTS

Transcriptomic analysis was performed by the Pôle Biomics of the Institut Pasteur Center for Technological Resources and Research (C2RT). We thank Dr. Philippe Dussart for providing the JEV SA14-14-2 vaccine, Dr. Yi-Lin Ling for providing the JEV-RP9 cDNA clone and Dr. Yoshiharu Matsuura for providing the anti-JEV NS5 antibody. This work was supported by a grant from the Seventh Framework Program (FP7) under grant number 278433-PREDEMICS. CK was funded by the French Ministry of Defense / Délégation Générale de l’Armement. MAD-S was funded by the DARPA INTERCEPT program (DARPA cooperative agreement #HR0011-17-2-0023. Please note that the content of the article does not necessarily reflect the position or the policy of the U.S. government and no official endorsement should be inferred).

## REFERENCES

1. Lindenbach BD, T.H., Rice CM., Flaviviridae: The viruses and their replication. Fields Virology, 5th ed. Philadelphia, PA. Lippincot-Raven Publishers., 2007: p. 1101–1152.

2. Campbell, G.L., et al., Estimated global incidence of Japanese encephalitis: a systematic review. Bull World Health Organ, 2011. 89(10): p. 766–74, 774A-774E.

3. Solomon, T., et al., Japanese encephalitis. J Neurol Neurosurg Psychiatry, 2000. 68(4): p. 405–15.

4. Yun, S.I. and Y.M. Lee, Japanese encephalitis: the virus and vaccines. Hum Vaccin Immunother, 2014. 10(2): p. 263–79.

5. Chambers, T.J., et al., Yellow fever/Japanese encephalitis chimeric viruses: construction and biological properties. J Virol, 1999. 73(4): p. 3095–101.

6. Eckels, K.H., et al., Japanese encephalitis virus live-attenuated vaccine, Chinese strain SA14-14-2; adaptation to primary canine kidney cell cultures and preparation of a vaccine for human use. Vaccine, 1988. 6(6): p. 513–8.

7. Liu, Y., et al., Safety of Japanese encephalitis live attenuated vaccination in post-marketing surveillance in Guangdong, China, 2005-2012. Vaccine, 2014. 32(15): p. 1768–73.

8. Eigen, M., Viral quasispecies. Sci Am, 1993. 269(1): p. 42–9.

9. Domingo, E., J. Sheldon, and C. Perales, Viral quasispecies evolution. Microbiol Mol Biol Rev, 2012. 76(2): p. 159–216.

10. Myint, K.S., et al., Neuropathogenesis of Japanese encephalitis in a primate model. PLoS Negl Trop Dis, 2014. 8(8): p. e2980.

11. Li, F., et al., Viral Infection of the Central Nervous System and Neuroinflammation Precede Blood-Brain Barrier Disruption during Japanese Encephalitis Virus Infection. J Virol, 2015. 89(10): p. 5602–14.

12. Chang, C.Y., et al., Disruption of in vitro endothelial barrier integrity by Japanese encephalitis virus-Infected astrocytes. Glia, 2015. 63(11): p. 1915–1932.

13. Chen, C.J., et al., Infection of pericytes in vitro by Japanese encephalitis virus disrupts the integrity of the endothelial barrier. J Virol, 2014. 88(2): p. 1150–61.

14. Liou, M.L. and C.Y. Hsu, Japanese encephalitis virus is transported across the cerebral blood vessels by endocytosis in mouse brain. Cell Tissue Res, 1998. 293(3): p. 389–94.

15. Tuma, P. and A.L. Hubbard, Transcytosis: crossing cellular barriers. Physiol Rev, 2003. 83(3): p. 871–932.

16. Yun, S.I., et al., Comparison of the live-attenuated Japanese encephalitis vaccine SA14-14-2 strain with its pre-attenuated virulent parent SA14 strain: similarities and differences in vitro and in vivo. J Gen Virol, 2016. 97(10): p. 2575–2591.

17. Yun, S.I., et al., A molecularly cloned, live-attenuated japanese encephalitis vaccine SA14-14-2 virus: a conserved single amino acid in the ij Hairpin of the Viral E glycoprotein determines neurovirulence in mice. PLoS Pathog, 2014. 10(7): p. e1004290.

18. Gromowski, G.D., C.Y. Firestone, and S.S. Whitehead, Genetic Determinants of Japanese Encephalitis Virus Vaccine Strain SA14-14-2 1. That Govern Attenuation of Virulence in Mice. J Virol, 2015. 89(12): p. 6328–37.

19. Melian, E.B., et al., NS1’ of flaviviruses in the Japanese encephalitis virus serogroup is a product of ribosomal frameshifting and plays a role in viral neuroinvasiveness. J Virol, 2010. 84(3): p. 1641–7.

20. Ye, Q., et al., A single nucleotide mutation in NS2A of Japanese encephalitis-live vaccine virus (SA14-14-2) ablates NS1’ formation and contributes to attenuation. J Gen Virol, 2012. 93(Pt 9): p. 1959–64.

21. Abbott, N.J., Blood-brain barrier structure and function and the challenges for CNS drug delivery. J Inherit Metab Dis, 2013. 36(3): p. 437–49.

22. Helms, H.C., et al., In vitro models of the blood-brain barrier: An overview of commonly used brain endothelial cell culture models and guidelines for their use. J Cereb Blood Flow Metab, 2016. 36(5): p. 862–90.

23. Weksler, B., I.A. Romero, and P.O. Couraud, The hCMEC/D3 cell line as a model of the human blood brain barrier. Fluids Barriers CNS, 2013. 10(1): p. 16.

24. da Costa, A., et al., Innovative in cellulo method as an alternative to in vivo neurovirulence test for the characterization and quality control of human live Yellow Fever virus vaccines: A pilot study. Biologicals, 2018. 53: p. 19–29.

25. da Costa, A., et al., A Human Blood-Brain Interface Model to Study Barrier Crossings by Pathogens or Medicines and Their Interactions with the Brain. J Vis Exp, 2019(146).

26. Liang, J.J., et al., A Japanese encephalitis virus vaccine candidate strain is attenuated by decreasing its interferon antagonistic ability. Vaccine, 2009. 27(21): p. 2746–54.

27. de Wispelaere, M., M.P. Frenkiel, and P. Despres, A Japanese encephalitis virus genotype 5 molecular clone is highly neuropathogenic in a mouse model: impact of the structural protein region on virulence. J Virol, 2015. 89(11): p. 5862–75.

28. Katoh, H., et al., Heterogeneous nuclear ribonucleoprotein A2 participates in the replication of Japanese encephalitis virus through an interaction with viral proteins and RNA. J Virol, 2011. 85(21): p. 10976–88.

29. Siflinger-Birnboim, A., et al., Molecular sieving characteristics of the cultured endothelial monolayer. J Cell Physiol, 1987. 132(1): p. 111–7.

30. Hsieh, J.T., et al., Japanese encephalitis virus neuropenetrance is driven by mast cell chymase. Nat Commun, 2019. 10(1): p. 706.

31. Turtle, L. and T. Solomon, Japanese encephalitis - the prospects for new treatments. Nat Rev Neurol, 2018. 14(5): p. 298–313.

32. Mustafa, Y.M., et al., Pathways Exploited by Flaviviruses to Counteract the Blood-Brain Barrier and Invade the Central Nervous System. Front Microbiol, 2019. 10: p. 525.

33. Liu, T.H., et al., The blood-brain barrier in the cerebrum is the initial site for the Japanese encephalitis virus entering the central nervous system. J Neurovirol, 2008. 14(6): p. 514–21.

34. Agrawal, T., et al., Japanese encephalitis virus disrupts cell-cell junctions and affects the epithelial permeability barrier functions. Plos One, 2013. 8(7): p. e69465.

35. Wang, K., et al., IP-10 Promotes Blood-Brain Barrier Damage by Inducing Tumor Necrosis Factor Alpha Production in Japanese Encephalitis. Front Immunol, 2018. 9: p. 1148.

36. Lai, C.Y., et al., Endothelial Japanese encephalitis virus infection enhances migration and adhesion of leukocytes to brain microvascular endothelia via MEK-dependent expression of ICAM1 and the CINC and RANTES chemokines. J Neurochem, 2012. 123(2): p. 250–61.

37. Cecchelli, R., et al., A stable and reproducible human blood-brain barrier model derived from hematopoietic stem cells. Plos One, 2014. 9(6): p. e99733.

38. Lippmann, E.S., et al., Derivation of blood-brain barrier endothelial cells from human pluripotent stem cells. Nat Biotechnol, 2012. 30(8): p. 783–1.

39. Doster, A., et al., Unfractionated Heparin Selectively Modulates the Expression of CXCL8, CCL2 and CCL5 in Endometrial Carcinoma Cells. Anticancer Res, 2016. 36(4): p. 1535–44.

40. Lin, X., et al., Insights into Human Astrocyte Response to H5N1 Infection by Microarray Analysis. Viruses, 2015. 7(5): p. 2618–40.

